# Antiretroviral APOBEC3 Cytidine Deaminases Alter HIV-1 Provirus Integration Site Profiles

**DOI:** 10.1101/833475

**Authors:** Hannah O. Ajoge, Tyler M. Renner, Kasandra Bélanger, Hinissan P. Kohio, Macon D. Coleman, Marc-André Langlois, Stephen D. Barr

## Abstract

APOBEC3 (A3) proteins are host-encoded deoxycytidine deaminases that provide an innate immune barrier to retroviral infection, notably against HIV-1. While the catalytic activity of these proteins can induce catastrophic hypermutation in proviral DNA leading to near-total restriction of infection, sublethal levels of deamination contribute to the genetic evolution of HIV-1. So far, little is known about how A3 might impact HIV-1 integrations into human chromosomal DNA. Using a deep sequencing approach, we analyzed the influence A3F and A3G on HIV-1 integration site selections. DNA editing was detected at the extremities of the long terminal repeat regions of the virus. Both catalytic active and non-catalytic A3 enzymes decreased insertions into gene coding sequences and increased integration sites into SINE elements, oncogenes and transcription-silencing non-B DNA features. Our data implicate A3 as host factors that influence HIV-1 integration site selection and promote insertions into genomic sites that are transcriptionally less active.

**GRAPHICAL ABSTRACT:** Schematic depicting the influence of APOBEC3 (A3) proteins on HIV integration site targeting.
*Left*, in the absence of A3, HIV has a strong preference for integrating into genes. *Right*, both catalytic active and non-catalytic A3 mutants decrease integration into genes and increase integration into SINE elements and in transcription-silencing non-B DNA features.

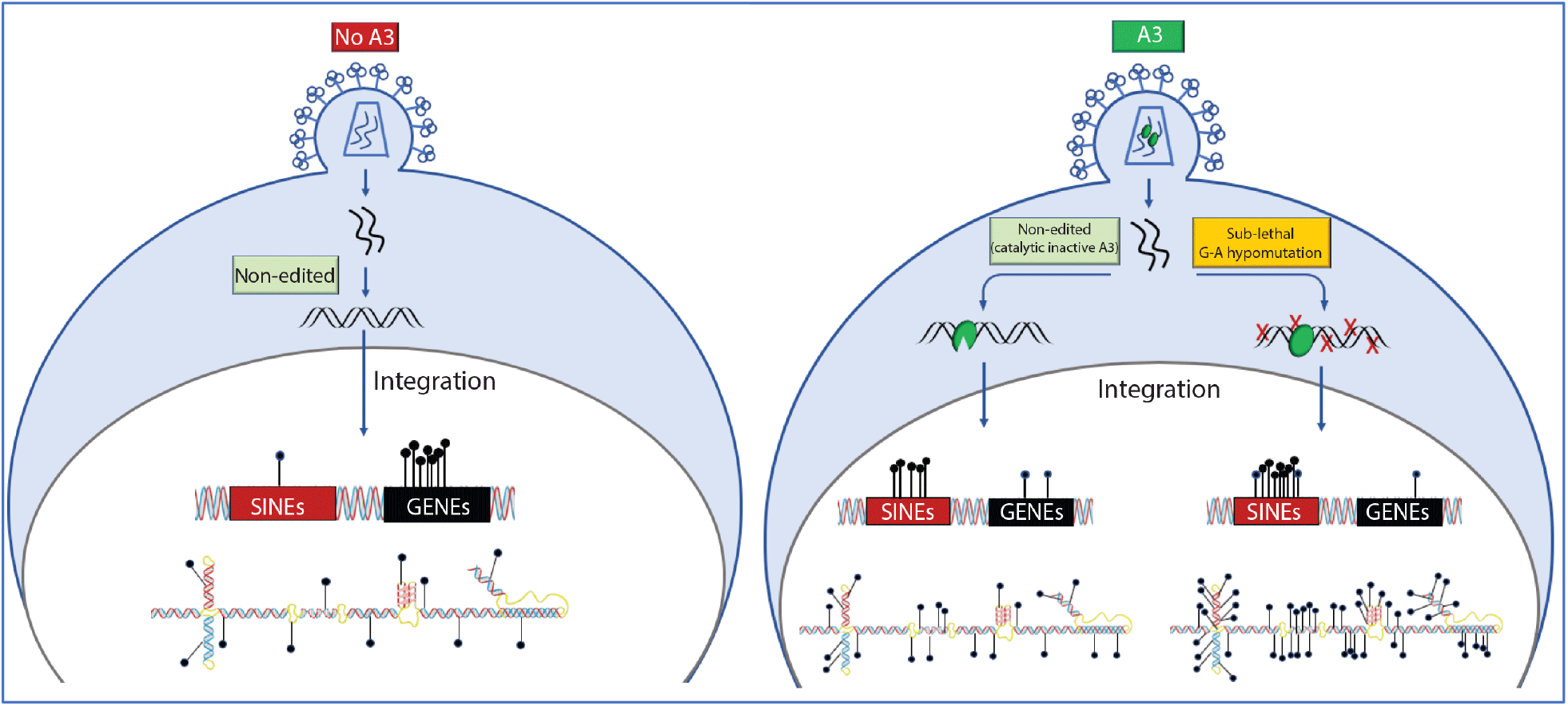

## INTRODUCTION

The human A3 family is comprised of seven members, five of which have demonstrated biologically relevant antiviral activity against HIV-1: APOBEC3D (A3D), APOBEC3F (A3F), APOBEC3G (A3G), certain haplotypes of APOBEC3H (A3H), and one polymorphic variant of APOBE3C (A3C)^1–4^. When HIV-1 infects a new CD4+ monocyte or lymphocyte, A3 proteins associate with viral proteins and RNA, resulting in their encapsidation within nascent egressing virions^5^. Virion-packaged A3 then exert their antiretroviral activity in the target cells during reverse transcription primarily by deaminating cytosines (C) into uracils (U) in negative sense single-stranded viral DNA (vDNA) replication intermediates^6–8^.

Very high levels of deamination, called hypermutation, are observed early in the infection that thoroughly inactivate the virus^6^. However, HIV-1 can overcome the effects of A3 proteins by the increased expression viral infectivity factor (Vif), which binds to and induces the polyubiquitination of the five anti-HIV A3 proteins, thereby orchestrating their progressive depletion by proteasomal degradation^9–11^. Consequently, nascent egressing virions package decreasing amounts of A3 proteins until the proteins have been expunged from the cytosol by Vif^12^. These viruses devoid of A3, or even with highly reduced levels of the restriction factor, can freely infect new cells to help rapidly spread the infection. Retaining low rates of A3 mutagenesis, or hypomutation, are believed to be important contributors to the rapid genetic evolution of HIV-1^13^.

A3 proteins can also restrict HIV-1 replication via mechanisms other than deamination (e.g. binding the viral RNA or to the viral reverse transcriptase (RT), which reduces vDNA synthesis)^14–20^. It was previously shown that A3G and A3F can interact with the viral integrase (IN) and RT, but the role of this binding on viral integration is not yet clear^21–24^. More importantly, it was shown that A3F and A3G proteins can compromise viral integration efficiency by modifying or altering adequate processing of the extremities of the long terminal repeats (LTR) of the virus^25,26^. It is unknown how this may affect HIV-1 proviral integration site selection.

Upon synthesis of proviral DNA, a pre-integration complex (PIC) comprised of viral and host (e.g. A3) proteins translocates to the nucleus in preparation for integration^27,28^. Proviral DNA integration into open chromatin involves host Lens Epithelium-Derived Growth Factor (LEDGF/p75) binding to the viral IN and polyadenylation specificity factor 6 (CPSF6) at the LTR ends (i.e. the intasome)^29–32^. This HIV-1 intasome favors integration in chromatin that is bent and associated with histones, active transcription units, regions of high G/C content, high gene density, high CpG island density, high frequencies of short interspersed nuclear elements (SINEs) (e.g. Alu repeats), epigenetic modifications and specific nuclear regions such as close to nuclear pore complexes^32–35^. In addition, non-B DNA structures potentially influence HIV-1 integration site targeting^36^. At least 10 non-B DNA conformations exist including A-phased motifs, inverted repeats, direct repeats, cruciform DNA, guanine quadruplex (G4) DNA, slipped DNA, mirror repeats, short-tandem repeats, triplex repeats and Z-DNA^37–40^.

While A3 proteins primarily act during reverse transcription to restrict HIV-1 through both deamination-dependent and independent mechanisms, A3F, and to a lesser extent A3G, remains associated with the PIC while it traffics into the nucleus^41^. In this study, we investigated the influence of A3 proteins on HIV-1 integration site selection. We found that both A3F and A3G have an important impact on integration site selection with both A3 deaminase-dependent and - independent activities contributing to this effect.

## RESULTS

### A3F and A3G strongly inhibit HIV-1 infection and integration in a dose-dependent manner

Using transfected 293T cells, we produced HIV-1 (NL4-3_ΔVif/ΔEnv-eGFP_) pseudotyped with vesicular stomatitis virus envelope glycoprotein (VSV-G) in the presence of wild type (wt) or deamination-defective mutant forms of A3F and A3G. We also included the A3G nucleic acid-binding defective mutants A3G [W94A] and A3G [W127A]^14^. Within the non-catalytic N-terminal A3G domain, W94 is part of the S<u>W</u>SPCxxC zinc-coordinating motif while W127 is within ARLYYF<u>W</u>. These tryptophan residues are important for general nucleic acid binding ability, substrate sequence recognition and protein oligomerization^14,21,42–44^. While both W94 and W127 mutations diminish RNA binding, the W127 substitution is unique because it prevents the homodimerization of A3G which results in a reduced processivity^14^. This highlights the importance of dimerization for the function of A3G catalytic activity in the presence of ssDNA^45^. Equal amounts of virus produced with each A3 were used to infect the permissive human T4-lymphoblastoid cell line CEM-SS. Productive infection of CEM-SS cells was determined forty-eight hours post-infection by flow cytometry by way of virus-encoded eGFP reporter expression (**Fig. 1A & S1**). Alternatively, infected cells were harvested for genomic DNA (gDNA) extraction for the quantification of proviral integration levels and the downstream analysis of integration sites. Three different amounts of input virus were used for the infections in addition to producing virus in the presence of increasing amounts of A3 proteins (**Fig. 1B & S1**). Increasing the amount of A3 plasmid used for virus production had a noticeable impact on HIV-1 particle release **(Fig. S2)**.

**Figure 1.**
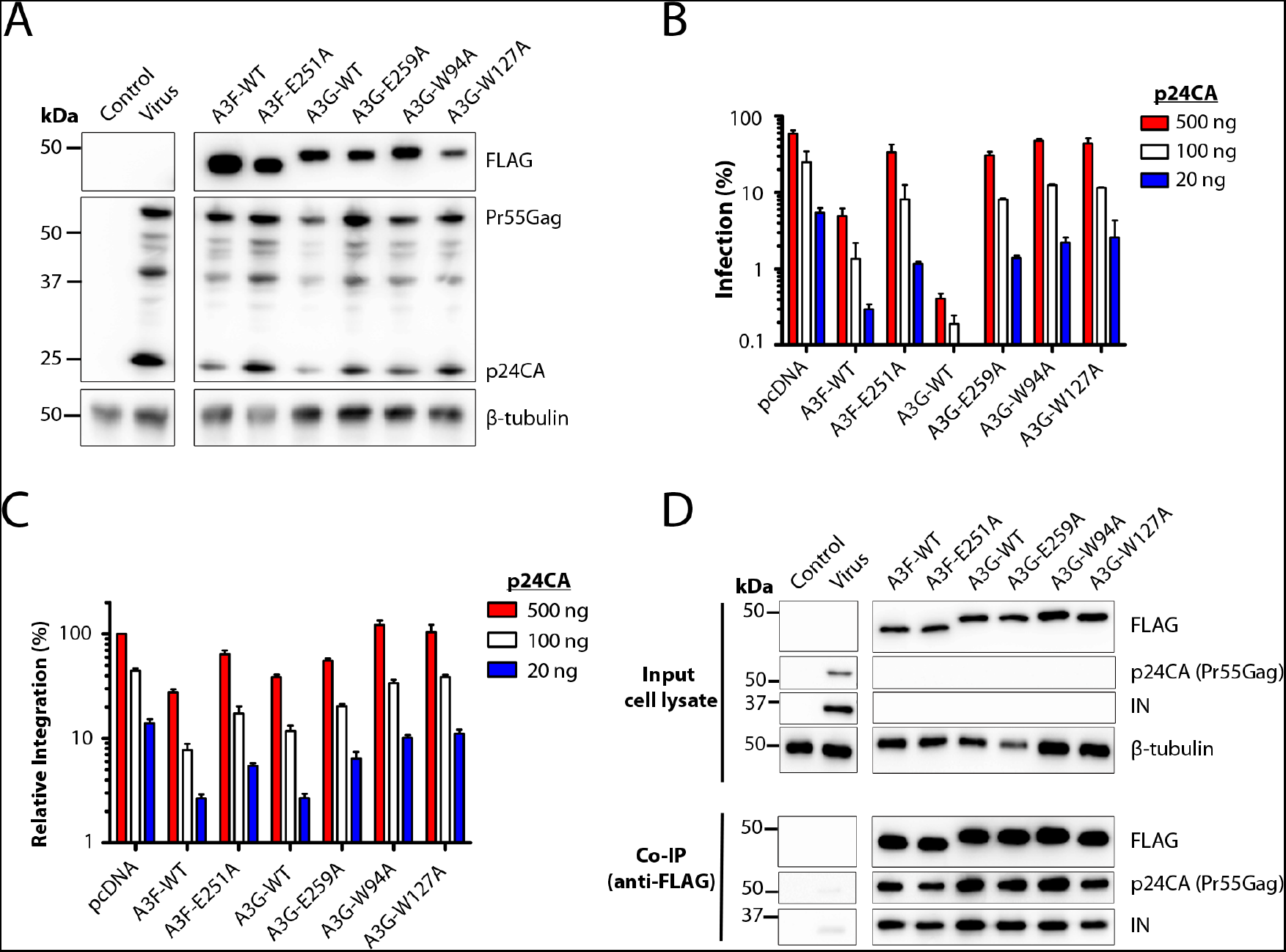
A3-mediated restriction of HIV-1 integration. **A**, Western blot analysis of virus producer cells. HIV-1 pseudotyped virus was produced in 293T cells by co-transfection of plasmids coding for NL4-3-ΔEnv/ΔVif/eGFP, VSV-G and either 100ng of empty pcDNA3 plasmid (Virus), or 100ng each of the A3 expression plasmids. The Control lane is the transfection of the pcDNA3 in absence of virus. Cell lysates were subjected to SDS-PAGE and Western blot analysis. **B**, CEM-SS cells were infected with the amounts of virus as indicated after normalization to capsid (p24CA) protein, as determined by ELISA. Infection was measured as the percentage of eGFP+ cells by flow cytometry. **C**, integrated provirus in the CEM-SS cells from (B) was quantified using Alu-based PCR combined with nested qPCR. **D**, 293T cells were transfected for the expression of each of the A3 proteins, HIV-1 (Virus) or pcDNA3 plasmid (Control). Viral and cell lysates were mixed together to improve detection and subjected to co-immunoprecipitation using an anti-FLAG antibody and analyzed by Western blotting. Data shown are representative of 3 independent experiments.

Potent restriction of HIV-1 was observed with wild type (wt) A3F, and even more notably with wt A3G (**Fig. 1B & S1**). Both catalytically inactive A3F [E251A] and A3G [E259A] demonstrated significantly less restriction. Our group has previously established that A3G [W94A] and A3G [W127A] each have diminished restriction capabilities but remained capable of viral DNA editing ^14^. These mutants had minor effects on the overall infection and integration efficiencies (**Fig. 1B and 1C**). Also, as expected, the level of inhibition by A3F and A3G were dependent on the amount of A3 proteins expressed during virus production (**Fig. S1**).

Given the multiple editing-dependent and -independent restriction mechanisms of A3, productive infection was correlated with overall levels of proviral integration. We measured relative integrated HIV-1 DNA by Alu-PCR followed by droplet digital PCR (ddPCR) targeting the HIV-1 LTR in one assay and the reporter eGFP gene as a control and input cellular DNA normalized to amplified actin DNA (**Fig. 1C & S3**)^46^. Integration levels tracked closely with infection levels, with the exception of the wt A3 proteins that both exhibit proportionately more apparent restriction of infection than integration. This is unsurprising, as eGFP reporter expression and fluorescence relies on the genetic integrity of its coding sequence which is frequently lethally mutated by A3F and A3G^47^.

### A3F and A3G interact with the viral Gag and IN

Several reports have shown that A3F and A3G interact with HIV-1 IN and Gag proteins, which are both components of the PIC^21–23,41,48,49^. However, binding to the A3 variants with inactive deaminase and defective nucleic acid binding have not been previously assessed in parallel. To characterize HIV-1 IN and Gag interactions with the various A3 proteins, lysates from cells expressing either FLAG-tagged A3 variants or HIV-1 were co-incubated and then coimmunoprecipitated using anti-FLAG and analyzed by Western blotting using anti-IN and anti-p24CA. As shown in **Fig. 1D**, all A3 proteins, including the deaminase-inactive and nucleic acid binding mutants co-immunoprecipitated with IN and Gag suggesting direct or indirect (i.e. complex) A3 interactions.

### A3F and A3G alter HIV-1 integration site targeting of genomic features

To identify HIV-1 integration sites, we amplified integration sites in gDNA isolated from cells infected with HIV-1 produced in the presence of the various A3 proteins. Integration site profiles were generated using the Barr Laboratory Integration Site Pipeline (BLISIP) as described^36,50^. BLISIP measures integration site enrichment in and near the genomic features CpG islands, DNAseI hypersensitivity sites (DHS), endogenous retroviruses, heterochromatic DNA regions (e.g. lamin associated domains (LADs) and satellite DNA), SINEs, long interspersed nuclear elements (LINEs), low complexity repeats (LCRs), oncogenes, genes, simple repeats and transcription start sites (TSS). In addition, BLISIP measures enrichment in and near the non-B DNA features A-phased motifs, cruciform motifs, direct repeats, G4 motifs, inverted repeats, mirror repeats, short tandem repeats, slipped motifs, triplex motifs and Z-DNA motifs.

Target cells infected with viruses containing A3F or A3G exhibited a significant increase in integration in and near SINEs compared to viruses not containing A3F or A3G (39% and 42% (respectively) compared to 17%; *P*<0.0001) (**Fig. 2A and 2B, Table S1 and Table S2**). Integration was also significantly enriched adjacent (1-500 nucleotides) to simple repeats for A3F and A3G in relation to the control with no A3 (26% and 35% (respectively) compared to 18%; *P*<0.05). In addition, integration with A3F and A3G was modestly increased in and near ERVs, LADs, oncogenes and LCRs compared to the no A3 control. Notably, in the presence of A3F and A3G, integration was significantly decreased in genes compared to the no A3 control (63% and 60% (respectively) compared to 75%; *P*<0.001). For comparison, 46% of integration sites occur in genes randomly (**Table S3**). Integration was also decreased in DHS and LINEs.

**Figure 2.**
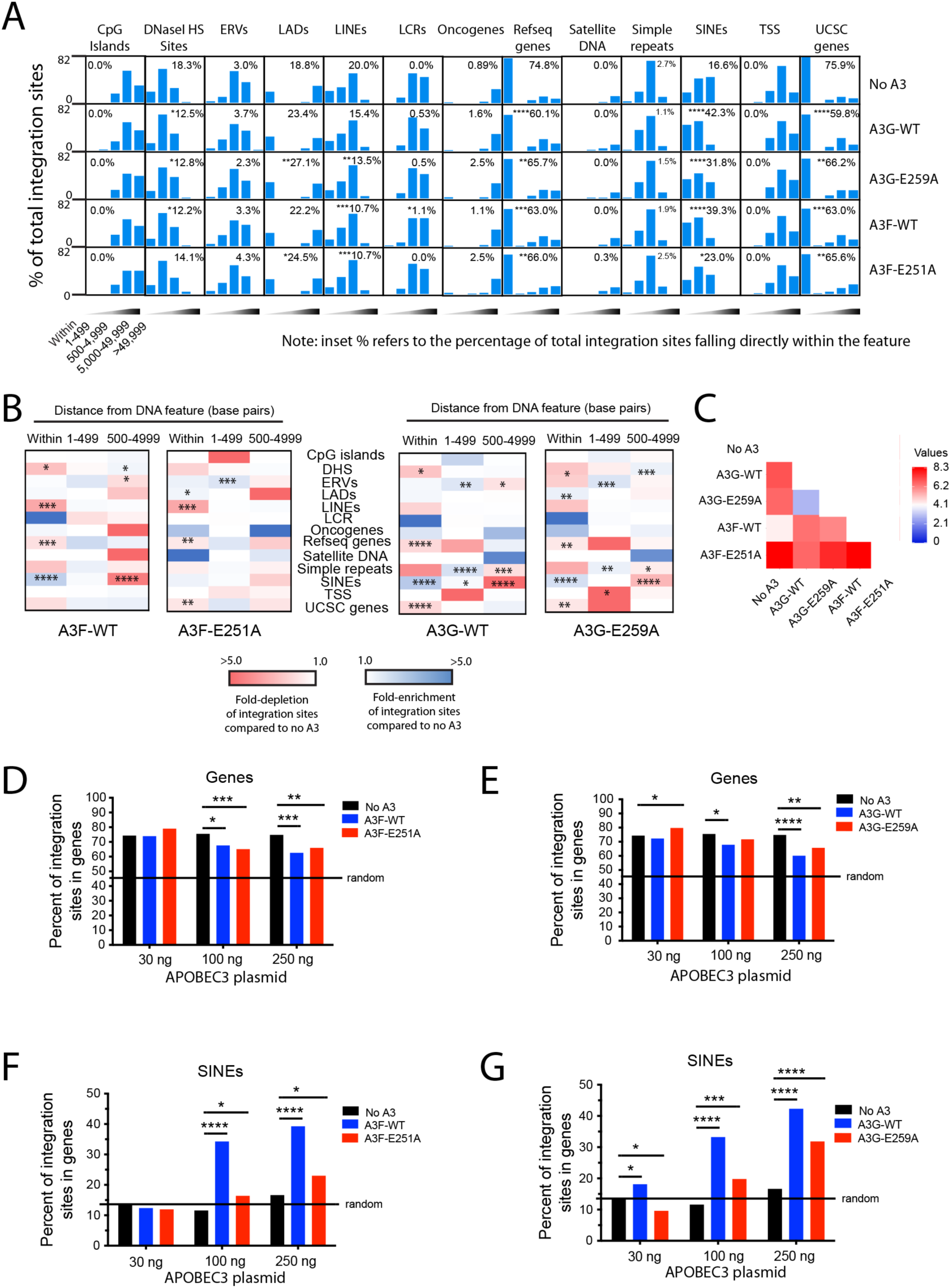
A3F and A3G expression alters integration site targeting of common genomic features. **A**, frequency of integration sites within or at different distance intervals (1-499 bp, 500-4999 bp, 5000-49999 bp or >49999 bp) away from various common genomic features in CEM-SS T cells infected with HIV-1 generated in the presence of A3F-WT, A3F [E251A], A3G-WT or A3G [E259A], or the absence of A3F or A3G (‘no A3’ control). **B**, heatmaps depicting the foldenrichment and depletion of integration sites at various distance intervals compared to the ‘no A3’ control virus. **C**, pairwise distance matrix was used to determine the overall similarity between the integration site profiles of CEM-SS cells infected with either the no A3 control virus or A3F-WT, A3F [E251A], A3G-WT or A3G [E259A] virus. The fold-enrichment and depletion values in each distance bin of each common DNA feature were used in the comparison. The heatmap shows the distance matrix calculated by Euclidean distance as the measurement method. Stronger relationships are indicated by the darker blue color and weaker relationships by darker red. **D** and **E**, percentage of total integration sites located in genes for CEM-SS cells infected with A3F-WT, A3F [E251A], A3G-WT or A3G [E259A] virus generated from cells expressing increasing concentrations of A3 protein. **F** and **G**, percentage of total integration sites located in SINES for CEM-SS cells infected with A3F-WT, A3F [E251A], A3G-WT or A3G [E259A] virus generated from cells expressing increasing concentrations of A3 protein. * *P* < 0.05, ** *P* < 0.01, *** *P* < 0.001, **** *P* < 0.0001; Fisher’s exact test.

Using the A3F and A3G mutants lacking deaminase activity, there was a significant increase in integration in SINEs and a significant decrease in integration in genes, but not to the same magnitude observed with their wt counterparts. Similar to their wild type counterparts, integration with A3F [E251A] and A3G [E259A] was modestly increased in ERVs, LADs and oncogenes (**Fig. 2A and 2B, Table S1 and Table S2**). To compare the overall similarity in integration site profiles between the various A3 constructs, we performed a pairwise analyses of the integration site profiles based on integration site enrichment or depletion within 5,000 bp of each genomic feature. As shown in **Fig. 2C**, the integration site profiles of the various A3-containing viruses differed substantially from each other, with A3G and A3G [E259A] sharing the most similarity in profiles.

Given that the largest differences in integration site selection were observed within genes and SINEs, we asked if these preferences were A3 dose-dependent. Indeed, increasing concentrations of A3 resulted in a decrease in the percentage of integrations sites in genes (**Fig. 2D and 2E and Table S3**). Conversely, increasing concentrations of A3 constructs resulted in a dose-dependent increase in the percentage of integrations sites in SINEs (**Fig. 2F and 2G and Table S3**). Together, these data show that A3F and A3G influence HIV-1 integration site targeting and that the deamination activity of A3F, and to a lesser extent A3G, influences the magnitude of this targeting.

### A3F and A3G expression alters integration site targeting of non-B DNA

We then determined the impact of A3 on targeting non-B DNA for integration. Cells infected in the presence of A3F or A3G exhibited enriched integration within 500 bp of most non-B DNA features (**Fig. 3A and Table S4**). Cells infected in the presence of A3F [E251A] or A3G [E259A] exhibited a similar level of integration near most non-B DNA compared to the control, except for direct repeats and slipped motifs, where a significant increase in integration was observed. Notably, A3F [E251A] exhibited a significant increase in integration near Z-DNA compared to the other A3 constructs and the control (25% compared to 10% for the control; *P*<0.0001).

**Figure 3.**
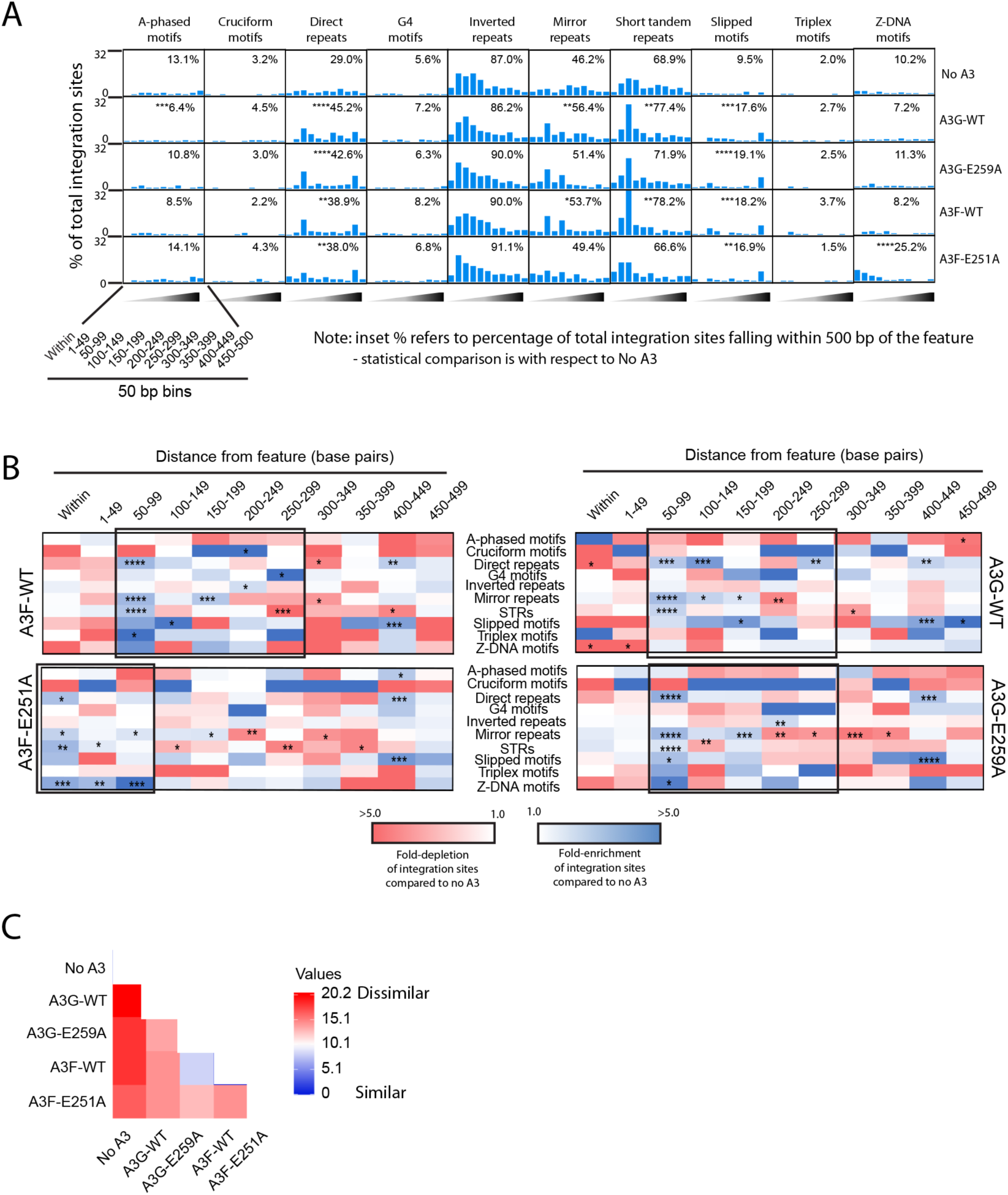
A3F and A3G alters integration site targeting of non-B DNA. **A**, frequency of integration sites within or in different 50 bp distance intervals (1-500 bp) away from various non-B DNA features in CEM-SS T cells infected with HIV-1 generated in the presence of A3F-WT, A3F [E251A], A3G-WT or A3G [E259A], or the absence of A3G or A3F (‘no A3’ control). **B**, heatmaps depicting the fold-enrichment and depletion of integration sites at various distance intervals compared to the ‘no A3’ control virus. Black boxes highlight regions of clustering. **C**, pairwise distance matrix was used to determine the overall similarity between the different integration site profiles. The fold-enrichment and depletion values in each distance bin for each non-B DNA feature were used in the comparison. The heatmap shows the distance matrix calculated by Euclidean distance as the measurement method. Stronger relationships are indicated by the darker blue color and weaker relationships by darker red. * *P* < 0.05, ** *P* < 0.01, *** *P* < 0.001, **** *P* < 0.0001; Fisher’s exact test.

To determine if there were differences in the distribution of integration sites near to the non-B DNA features, we compared the number of integration sites in bins of 50 bp up to 500 bp away from each non-B DNA motif (**Fig. 3B and Table S5**). Integration preferences of viruses produced with A3F, A3G and A3G [E259A] clustered in a region 50-300 bp away from the non-B DNA. A3F [E251A] differed from the others in that integration clustered in a region within 100 bp of the features. Pairwise analysis of the integration site profiles (within 500 bp of the features) showed that while A3F and A3G [E259A] shared a surprising amount of similarity, all other integration profiles are highly unique (**Fig. 3C**). Together, these data show that A3F and A3G influence HIV-1 integration site targeting of non-B DNA features with a substantial contribution of their deamination activities in the targeting of A-phased, mirror repeats, STRs and Z-DNA features.

### A3G residues W94 and W127 impact HIV-1 integration site targeting

We analyzed the integration site profiles of cells infected with virus produced in the presence of A3G [W94A] or A3G [W127A] mutants to determine if these residues impact the ability of wt A3G to influence integration site targeting. Compared to their wt counterpart, A3G [W94A] and A3G [W127A] exhibited a significant increase in integration in genes and decreased integration in SINEs (**Fig. 4A and 4B, and Tables S1 and S2**). In addition, A3G [W127A] exhibited a notable increase in integration in oncogenes compared to wt A3G. Interestingly, while A3G [W94A] exhibited an intermediate phenotype between the control and wt A3G, A3G [W127A] seemed to exacerbate the integration site preferences of HIV-1. Pairwise analyses of all A3 integration site profiles (within 5,000 bp of the various features) showed that A3G [W94A] was most similar to wt A3F and that A3G [W127] differed substantially from A3 variants tested (**Fig. 4C**).

**Figure 4.**
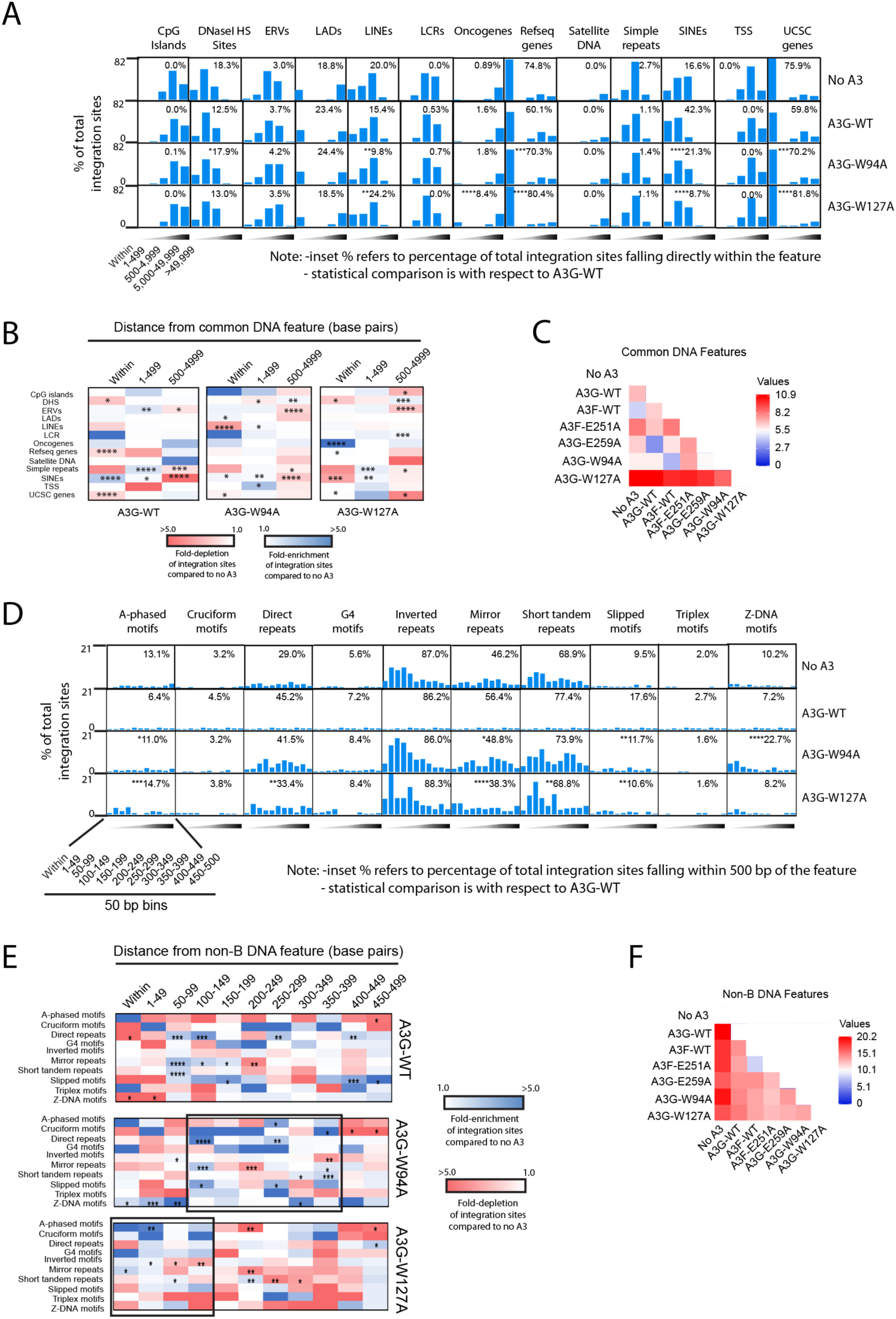
A3 residues W94 and W127 play a role in integration site targeting. **A**, frequency of integration sites within or at different distance intervals away from various common genomic features in CEM-SS T cells infected with HIV-1 generated in the presence of A3G-WT, A3G [W94A], A3G [W127A], or the absence of A3F or A3G (‘no A3’ control). **B**, heatmaps depicting the fold-enrichment and depletion of integration sites at various distance intervals compared to the ‘no A3’ control virus. **C**, pairwise distance matrix was used to determine the overall similarity between the integration site profiles of CEM-SS cells infected with either the no A3 control virus or A3F-WT, A3F [E251A], A3G-WT, A3G [E259A], A3G [W94A] or A3G [W127A] virus. The fold-enrichment and depletion values in each distance bin of each common DNA feature were used in the comparison. The heatmap shows the distance matrix calculated by Euclidean distance as the measurement method. Stronger relationships are indicated by the darker blue color and weaker relationships by darker red. **D**, frequency of integration sites within or at different distance intervals away from various non-B DNA features. **E**, heatmaps depicting the fold-enrichment and depletion of integration sites at various distance intervals compared to the ‘no A3’ control virus. Black boxes highlight regions of clustering. **F**, pairwise distance matrix was used to determine the overall similarity between the integration site profiles. * *P* < 0.05, ** *P* < 0.01, *** *P* < 0.001, **** *P* < 0.0001; Fisher’s exact test.

With respect to non-B DNA features, cells infected with A3G [W94A]- or A3G [W127A]-containing virus exhibited a significant increase in integration near A-phased motifs and decreased integration near mirror repeats and slipped motifs compared to wt A3G (**Fig. 4D and Table S4**). In addition, A3G [W127A] exhibited a significant increase in integration near Z-DNA. The distribution of integration sites in bins of 50 bp up to 500 bp away from each non-B DNA motif was similar between wt A3G and A3G [W94A], where sites clustered predominantly in a region 100-400 bp away from the features (**Fig. 4E and Table S5**). An exception was Z-DNA where sites were highly enriched in and within 100 bp of Z-DNA motifs. In contrast, integration sites from cells generated in the presence of A3G [W127A] tended to cluster within 150 bp of non-B DNA. Together, these data show that A3G residues W94, and to a greater extent W127, differentially impact the ability of A3G to influence integration site targeting.

### A3F and A3G reduce the number of hotspots and clustering of integration sites

The concept of an HIV-1 integration “hotspot” was introduced to describe areas of the genome where integrations accumulate more than expected by chance in the absence of any selection process^51^. Given our findings that A3F and A3G influence integration site targeting, we asked if they also impact the number of integration hotspots and clustering of sites. We defined an integration hotspot as a 1 kilobase (kb) gDNA fragment containing four or more unique integration sites. CEM-SS T cells infected with HIV-1 produced in the presence of A3F or A3G exhibited a substantial reduction in the number of hotspots compared to cells expressing no A3 (*P* < 0.05, Fisher’s exact test) (**Fig. 5A and Table S6**). A3F [E251A] and A3G [E259A] also exhibited a reduced number of hotspots indicating that the A3 deamination activities were not essential for this effect. In contrast, the presence of A3G [W94A] or A3G [W127A] exhibited no significant reduction in the number of integration site hotspots.

**Figure 5.**
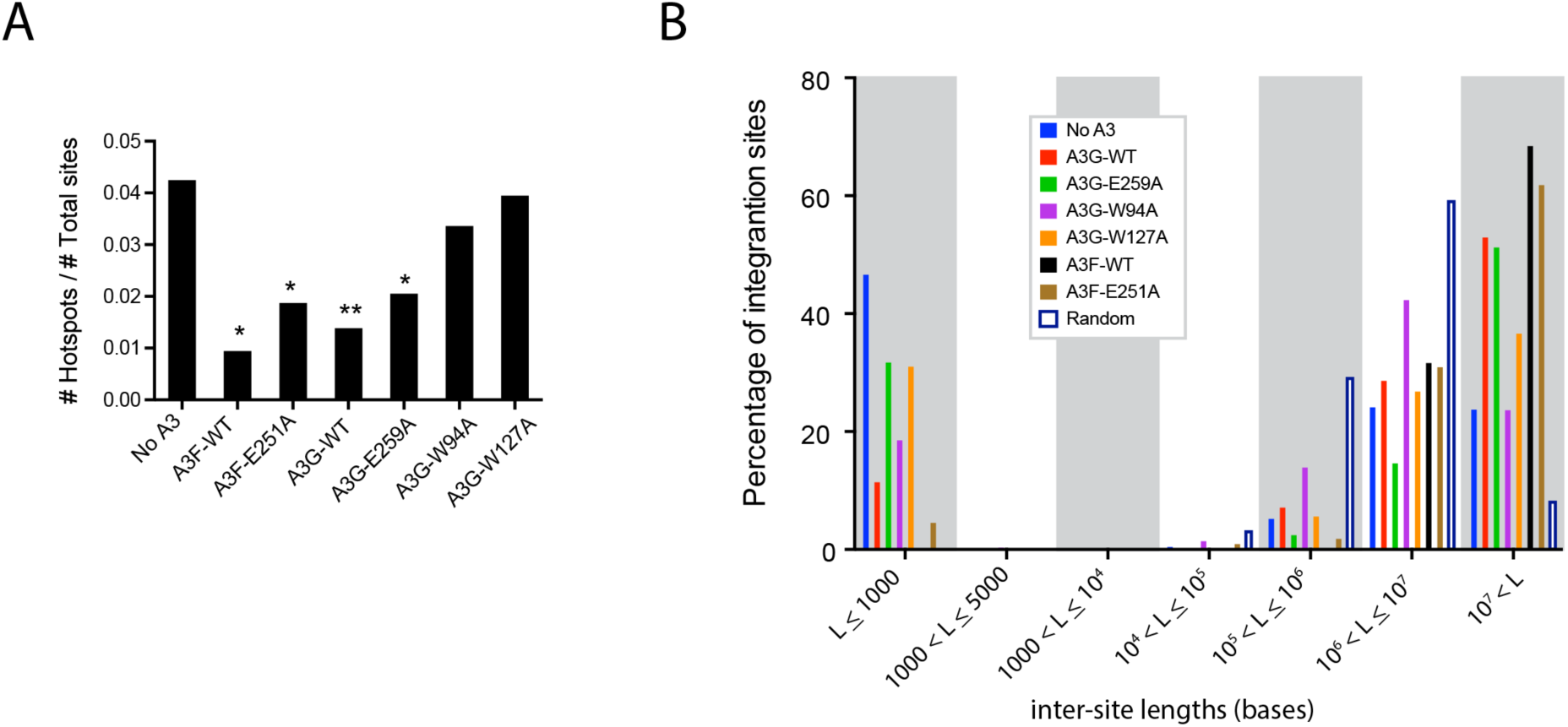
A3F and A3G reduce the number of integration hotspots and clustering of sites. **A**, Analysis of integration site hotspots. A hotspot was defined as a 1 kb window in the genome hosting 4 or more unique integration sites. Integration hotspots are shown as a proportion of total integration sites from CEM-SS cells infected with HIV-1 produced in the presence of no A3F or A3G (“no A3 control”), or from cells expressing A3F-WT, A3F [E251A], A3G-WT, A3G [E259A], A3G [W94A] or A3G [W127A]. **B**, Analysis of integration site spacing. Nucleotide distances between sites were measured and collected in length (L) bins with the shortest lengths to the left and longest to the right. A matched random control dataset was generated *in silico*. * *P* < 0.05, ** *P* < 0.01; Fisher’s exact test.

To document clustering of integration sites within genomic regions, we compared the distance between proviral integration sites (**Fig. 5B**). The control population of integration sites contained more short intersegment distances than expected by chance (*i.e*. random distribution), indicative of clustering. Viruses produced with A3F or A3G exhibited a striking reduction in clustering compared to the control cells (*P* <0.0001, Fisher’s exact test). This reduction was lost in virus containing A3G [E259A] and A3G [W127A], and to a lesser extent A3G [W94A] and A3F [E251A]. Together, these data show that expression of A3F and A3G reduce the number of integration hotspots and clustering of integration sites.

### G-to-A mutations in the LTR alter integration site targeting

To determine if deamination of LTR ends impacted integration site targeting, we aligned unique integrated 3’ LTR nucleotide sequences and represented them graphically as sequence logos using WebLogo^52,53^ (**Fig. 6A**). As expected, the 3’ LTR ends of the control and the deamination-defective A3 mutants A3F [E251A] and A3G [E259A] were highly similar. The 3’ LTR ends of A3F and A3G were also highly similar with the exception of the 2 nucleotides located at positions 14 and 15 from the end of the LTR. In the presence of A3F, 80% of the LTRs contained GG at these positions, 16% contained GA, 1% contained AG and 2% contained AA (**Fig. 6B**). In the presence of A3G, 67% of the LTRs contained GG at these positions, 1% contained GA, 30% contained AG and 2% contained AA.

**Figure 6.**
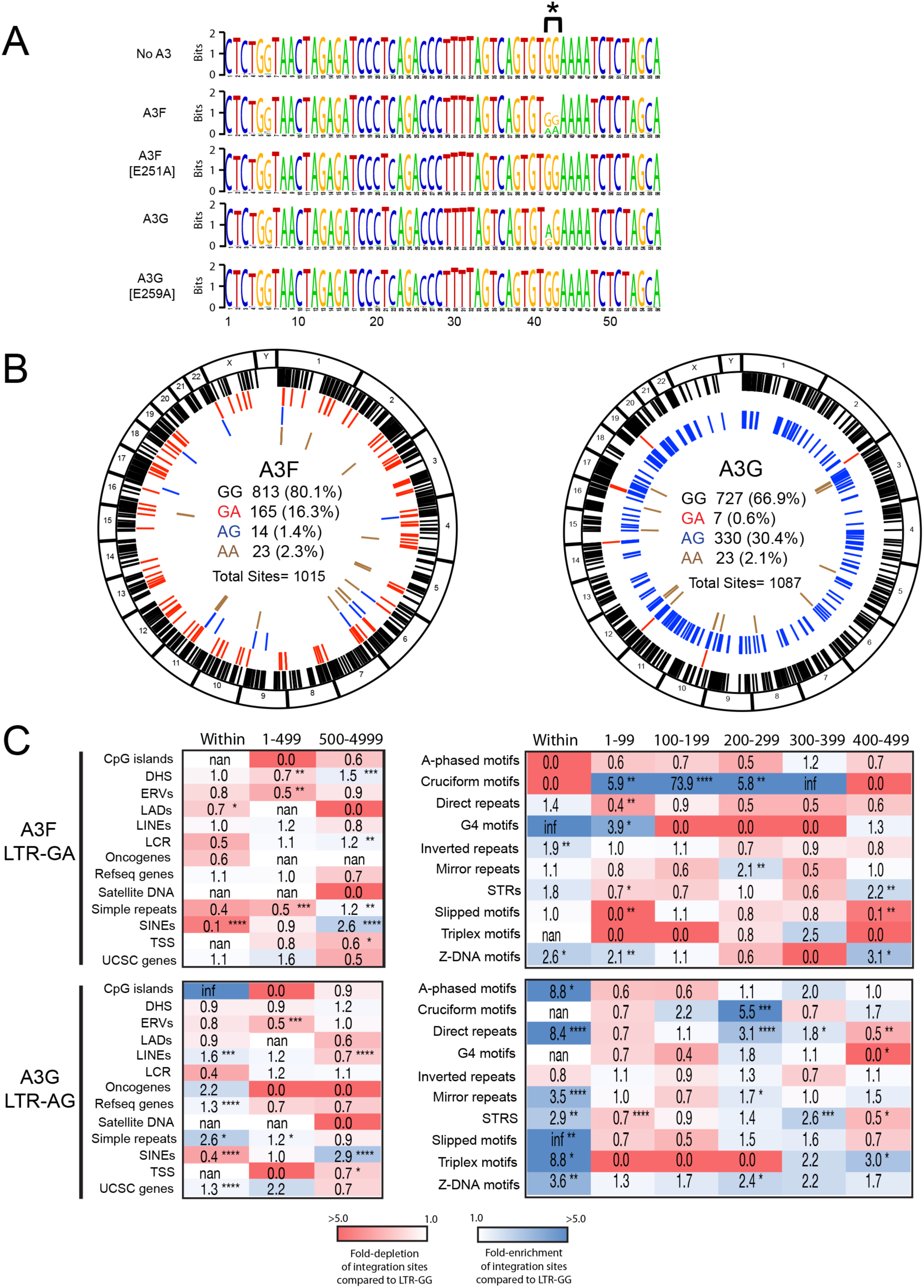
G to A mutations in the 3’ LTR alters integration site targeting. **A**, LOGO representations of the terminal 56 nucleotides of the 3’ LTRs of all integrated HIV-1 proviruses generated in the presence of either A3F, A3F [E251A], A3G, or A3G [E259A]. **B**, Circa plots showing the integration sites of A3F (left) or A3G (right) proviruses in the genome of infected CEM-SS cells. The outer ring represents the different chromosomes. The locations of integration sites of proviruses containing GG (black), GA (red), AG (blue) and AA (brown) at positions 14 and 15 nucleotides from the LTR end are represented as colored ticks. The number and percentage of total sites are shown inside the circa plots. **C**, heatmaps depicting the fold-enrichment and depletion of integration sites at various distance intervals from common genomic features (left) and non-B DNA features (right). Integration sites from A3F LTR-GA (top) and A3G LTR-AG (bottom) proviruses are shown. Fold changes are with respect to A3F LTR-GG and A3G LTR-GG respectively. * *P* < 0.05, ** *P* < 0.01, *** *P* < 0.001, **** *P* < 0.0001; Fisher’s exact test. Infinite number (inf) represents 1 or more integrations were observed when 0 integrations were expected by chance. Not a number (nan) represents 0 integrations were observed and 0 were expected by chance.

To determine if G-to-A mutations at positions 14 and/or 15 from the 3’ LTR end correlated with an altered integration site profile, we compared the A3F and A3G integration site profiles of proviruses with LTR ends containing either GA or AG (respectively) to that with GG at these positions. A3F LTRs containing the GA dinucleotide (A3F-LTR-GA) exhibited a notable reduction in integration sites within SINEs and increased integration more distal (500-5000 bp) to SINEs (**Fig. 6C**). Increased integration was also observed more distal to DHS, LCR and simple repeats. Strikingly, the integration site profile changed dramatically with respect to non-B DNA features. A3F-LTR-GA sites were highly enriched (up to 74-fold) 1-400 bp from cruciform motifs (**Fig. 6C**). Additionally, sites were enriched in and near G4 DNA, inverted repeats and Z-DNA. A3G LTRs containing the AG dinucleotide (A3G-LTR-AG) also exhibited reduced integration in SINEs and increased integration more distal to SINEs; however, unlike A3F-LTR-GA, integration was also increased in genes and simple repeats. Similar to A3F-LTR-GA, A3G-LTR-AG exhibited a striking change in integration site profile with respect to non-B DNA. Integration was enriched in and/or near most non-B DNA features. Together, these data show that A3-induced G-to-A mutations at position 14 or 15 from the end of the 3’ LTR correlates with a significantly altered integration site profile.

## DISCUSSION

Despite the presence of numerous cellular host restriction factors that collaboratively work to inhibit early stages of HIV-1 infection, integration of the HIV-1 genome into the host genome still occurs. This integration event will have varying consequences depending on the genomic site of integration. These may involve direct modulation of host gene networks, controlling levels of viral transcription, and in some cases influence the outcome of active or latent infection^54,55^. Normally, A3 proteins play a major role in hindering reverse transcription of the HIV-1 RNA genome; however, HIV-1 Vif circumvents this restriction by reducing A3 protein levels. Despite this, some A3 proteins including A3F and A3G have been shown to interact with the PIC and translocate into the nucleus^41^. The consequences of this localization in the nucleus and its impact on HIV-1 integration was previously unknown. Here we have shown that A3F and A3G significantly alter HIV-1 integration site selection.

HIV-1 has an integration site preference for actively transcribing genic regions, particularly those activated during the infection^56,57^. The selection process undoubtedly plays an important role in expansion and persistence of infected cells, as demonstrated in patients on cART^58^. Host cellular proteins are known to play critical roles in HIV-1 integration site selection. For example, LEDGF/p75 and CPSF6 promote integration into actively transcribing genes residing in gene-dense regions^29,30,32,57,59–63^. Here we showed that host cellular A3 expression shifts the accumulation of HIV-1 integration sites away from genes and towards SINEs in a dose-dependent manner. Moreover, increased A3 expression reduced the number of integration site hotspots in the human genome suggesting that A3 increases integration site diversity. Given the ability of A3 to bind integrase within the PIC, it will be interesting to learn if A3 interferes with the ability of integration site targeting factors such as LEDGF and CPSF6 to bind and target the PIC to transcriptionally active regions of the nucleus and genome. Of course, during later rounds of HIV-1 replication, A3 levels are reduced by Vif, which could be another mechanism by which the virus promotes integration into more active regions of the genome to help establish productive infection. Given this temporal gradient of A3 expression, their effects on integration likely occur at the earlier moments after a new cell is infected when it releases permissive levels of A3 in HIV virions. Additionally, there is evidence that infectious deaminated proviral genomes do exist, and therefore support an opportunity for A3 to influence integration^64–67^. While most of these deaminated proviral genomes are defective, some can still produce HIV RNA, viral protein and thereby contribute to chronic immune activation in absence of virus egress^55^. Since integration sites also play a critical role in the expansion and persistence of HIV-1-infected cells, these A3-directed integration sites could be important in the context of chronic infection. More work is needed to determine the contribution, if any, of A3 to such long-lived infected cells.

The ability of A3 to alter the integration site profile was partially dependent upon the deaminase activity of both A3F and A3G, but more strongly dependent upon the nucleic acid binding ability of A3G. We previously showed that the nucleic acid binding mutants A3G [W94A] and A3G [W127A] are encapsidated into virus particles, albeit to a reduced degree, and exhibited similar deaminase activity compared to wt A3G^14^. In addition, these mutants did not reduce late reverse transcripts or integration of the provirus^14^. Our finding that these same mutants displayed unique influences on the integration site selection of HIV-1 PICs when compared to the other A3 constructs may imply that deaminase-independent activity is an important factor, but not the sole factor, in influencing integration site selection. A key difference between the A3G [W94A] and A3G [W127A] mutants is that while A3G [W94A] can form homodimers, the A3G [W127A] mutant is less proficient in this regard^14^. Thus, the differences observed between A3G and A3G [W94A] may be due to a reduced affinity for nucleic acids, whereas the differences observed between A3G [W94A] and A3G [W127A] may be due to multimerization defects^14^.

Genomic position effects have been shown to influence HIV-1 expression and latency reversal^68–70^. LADs represent a repressive chromatin environment tightly associated with the nuclear periphery^68,71^. SINEs (e.g. Alu repeats) and other transposed sequences are known to serve as direct silencers of gene expression due to their repressed chromatin marks (histone H3 methylated at Lys9)^72–74^. Moreover, some non-B DNA structures including G4, cruciform, triplex and Z-DNA have been shown to potently silence expression of adjacent genes^75–85^. We showed here that both A3F and A3G increased the frequency of HIV-1 integration in or near LADs, SINEs and several non-B DNA structures, potentially implicating A3 in promoting integration in more transcriptionally silent regions of the genome. Interestingly, proviruses with deaminated 3’ LTR ends were highly enriched in and near gene silencing non-B DNA structures compared to proviruses with non-deaminated 3’ LTR ends. It is currently unknown if these proviruses are replication-competent, or if this integration site bias is a direct impact of deamination and/or directed by A3 itself. Further work is required to better understand this integration site bias and its impact, if any, on proviral silencing.

In conclusion, we have shown for the first time that both deaminase-active and -inactive A3F and A3G enzymes can bind the viral IN and alter the integration site profiles of HIV-1. Currently, the overall functional impact of A3 on the integration site profiles of HIV-1 and disease progression is unclear and will require additional research. A3 interfering with integration may represent a last-ditch effort to direct the intasome away from active genes and into more transcriptionally silent regions of the genome to promote proviral silencing. However, the unintended consequences of such actions may be to contribute to the latent reservoir which remains the greatest hurdle to overcome if a cure for HIV is to be developed.

## MATERIALS AND METHODS

### Cell lines and plasmids

Cell lines were maintained in complete media (10% FBS, 100U/mL Penicillin and 100 μg/mL Streptomycin). HEK 293T cells (ATCC CRL-3216™) were maintained in complete DMEM with high glucose. CEM-SS cells (NIH AIDS #776) were maintained in complete RPMI. Both cell stocks were maintained in a humidified 37°C incubator with 5% CO_2_. NL4-3-ΔVif/ΔEnv-eGFP was developed through site-directed mutagenesis of the NL4-3 ΔEnv-eGFP, which was originally obtained from the NIH AIDS Reagents Program (N.A.R.P.) (Catalog #11100)^14^. NL4-3-ΔVif/ ΔEnv-eGFP was pseudotyped with Vesicular Stomatitis Virus-G protein (VSV-G) (pMDG) as previously described^21,86^. pcDNA 3.1 (pcDNA3; Invitrogen) was used as an empty vector control for transfection and all A3 expressing plasmids have been described previously^21,86^.

### Virus production and infection

HEK 293T cells were seeded at 7.5 × 10^5^ cells in each well of a 6-well plate. 24 hours post-seeding, the cells were co-transfected with plasmids carrying the NL4-3-ΔEnv/ΔVif/eGFP reporter vector and pMDG, together with either empty vector or A3 plasmids as indicated using polyethylenimine (PEI)^87^. While the ratios of the NL4-3-ΔEnv/ΔVif/eGFP and pMDG plasmids remained constant (750 ng: 250 ng), the levels of co-transfected empty vector or A3 plasmid varied according to the experiment (30 ng, 100 ng and 250 ng). The total amount of DNA transfected was kept constant using empty vector (pcDNA3). Cells were incubated for 72 hours to produce virus. Virus production was confirmed using Western blotting with anti-p24Capsid (N.A.R.P.), anti-FLAG (Clone M2; Sigma), anti-β-Tubulin (ab21058; Abcam). Virus supernatants were collected, centrifuged at 500 x g for 5 minutes and filtered using a 450 nm syringe-filter to remove cellular debris. At this point, a Sandwich-ELISA was performed to determine levels of capsid protein (p24) using antibody isolated from Hybridoma 31-90-25 (#HB-9725; ATCC) and 183-H12-5C (N.A.R.P. #1513). Twenty hours after virus collection, CEM-SS cells were seeded in a 12-well plate at a density of 5 × 10^5^ cells per well and infected with normalized capsid levels (500 ng, 100 ng or 20 ng) by spinoculation for 1 hour at 900 x g without polybrene. Cells were incubated for 48 hours and collected for downstream flow cytometry analysis and gDNA extraction. Wizard gDNA Purification Kit (Promega) was used to isolate and purify gDNA from CEM-SS cells.

### Quantification of integrated provirus using Alu-based qPCR

PowerUp™ SYBR Master Mix (ThermoFisher) was used to quantify the relative levels of cells using a Viia™7 Real-Time PCR Instrument (Applied Biosystems) with 50 ng gDNA template using the following primers: Actin-FWD 5’-CAT GTA CGT TGC TAT CCA GGC-3’ and Actin-REV 5’-CTC CTT AAT GTC ACG CAC GAT-3’. Cycling conditions: initial denaturation at 95°C for 3 minutes, followed by 45 cycles of 95°C for 15 seconds and 60°C for 1 minute. Data was analyzed using the QuantStudio software. Next, similarly to a previously described protocol, Alu-PCR was performed using 50 ng of gDNA and PrimeStar GXL DNA Polymerase (Takara) using the following conditions: initial denaturation at 94 °C for 1 minute followed by 30 cycles of 98 °C for 10 seconds, 55 °C for 15 seconds and 68 °C for 10 minutes, ending with an additional extension step of 68 °C for 10 minutes^46^. All primers targeting the HIV-1 sequence were designed to exclude A3 dinucleotide hotspots to avoid inducing PCR biases. To quantify integrated eGFP sequences, the following primers were used: Alu1 5’-TCC CAG CTA CTG GGG AGG CTG AGG-3’, Alu2 5’-GCC TCC CAA AGT GCT GGG ATT ACA G-3’ and Lambda-eGFP-FWD 5’-ATG CCA CGT AAG CGA AAC TGT ACA ACT ACA ACA GCC ACA ACG TCT ATA TC-3’. A dilution of this was analyzed by ddPCR using the QX200 system (BioRad) with the following conditions: initial denaturation at 95 °C for 10 minutes followed by 45 cycles of 94 °C for 30 seconds, 60 °C for 30 seconds and 72 °C for 30 seconds. This was followed by a final denaturation of 98 °C for 10 minutes. The following primers were used: LambdaE-F2 5’-ATG CCA CGT AAG CGA AAC TGT ACA ACT AC-3’, HIV eGFP REV 5’-TGA GGA TTG CTT AAA GAT TAT TGT TTT ATT ATT T-3’. This probe was used: /5HEX/ CCC CGT GCT /ZEN/ GCT GCC CRA CAA CCA CTA CC /3IABkFQ. To quantify integrated 5’ LTR sequences, the following primers were used: Lambda-R-U5-REV1 5’-AGT TTC GCT TAC GTG GCA TCA GAC GGG CAC ACA CTA CTT TGA GCA C-3’, Alu1 Comp 5’-CCT CAG CCT CCC CAG TAG CTG GGA-3’ and Alu2 Comp 5’-CTGT AAT CCC AGC ACT TTG GGA GGC-3’. A dilution of this was analyzed by ddPCR as described using the same conditions described above and the following primers: LambdaR-REV2 5’-GTT TCG CTT ACG TGG CAT CAG ACG G-3’ and Late U3-FWD 5’-GCT ACA TAT AAG CAG CTG CTT TTT GCC TGT AC-3’. The following probe was used: /5YAkYel/ CTT TAT TGA GGC T+T AAG +C+ AG+ T+G +GG T/3IABkFQ. Nucleotides followed by a + (N+) indicate an LNA base to improve the melting temperature of the probe. Results from the Alu-PCR that quantified integrated proviruses using the eGFP sequence or the 5’ LTR sequence primers were then averaged.

### Immunoprecipitation

HEK 293T cells were transfected with NL4-3-ΔEnv/ΔVif/eGFP and VSV-G and the viral supernatant was cleared of cellular debris. Virus supernatant was concentrated by ultracentrifugation through a 20% sucrose cushion at 100,000 x g for 3 hours at 4 °C using a Type 70Ti. The viral pellet was resuspended in an isotonic 1% Triton-X 100 lysis buffer. At the same time, the virus producer cells were lysed using a soft lysis buffer with 500 mM NaCl to efficiently burst the nucleus and maximally release integrase. Protease inhibitors (Roche) were used at all times to prevent protein degradation. Salt-Active Nuclease (Sigma) was used to remove the gDNA according to the manufacturer’s protocol. Remaining cellular debris was removed by centrifugation at 4 °C at 17,000 x g for 10 minutes. Cellular and supernatant lysates were mixed together to maximize levels of viral components isolated. Overall salt levels were brought back to an isotonic state using sterile water. The aforementioned A3 or pcDNA 3.1 plasmids were each transfected individually in their own well at the same time as the NL4-3-ΔEnv/ΔVif/eGFP transfection. Seventy-two hours post-transfection, the cells were collected and lysed using an isotonic 1% Triton-X 100 lysis buffer. Lysates were sonicated to improve protein solubilization and centrifuged at 17,000 x g for 10 minutes at 4 °C to remove remaining cellular debris. A sample of each lysate was collected prior to immunoprecipitation to assess input levels. The viral lysate was equally divided among the cellular lysates containing A3 or controls. These lysates were then mixed with 30 μL of anti-FLAG conjugated magnetic μbeads (Miltenyi) and incubated on a tube rotator for 3 hours at 4 °C. The μbeads were then magnetically isolated using a μcolumn according to manufacturer’s instructions. Samples were denatured and analyzed by Western blotting using anti-IN (IN-2), anti-p24 Capsid (N.A.R.P.), anti-FLAG (Clone M2; Sigma), anti-β-Tubulin (ab21058; Abcam).

### Integration site library and computational analysis

Integration sites from three independent infections carried out with each A3 transfection condition (*i.e.*, 30ng, 100ng, and 250ng) and for each of the three amounts of virus used for the infection (*i.e.*, 20ng, 100ng and 500ng of p24) were pooled together accordingly.

Genomic DNA was processed for integration site analysis and sequenced using the Illumina MiSeq platform as described^36,50^. Fastq sequencing reads were quality trimmed and unique integration sites identified using our in-house bioinformatics pipeline^36^, which is now called the Barr Lab Integration Site Identification Pipeline (BLISIP version 2.9) and includes the following updates: bedtools (v2.25.0), bioawk (awk version 20110810), bowtie2 (version 2.3.4.1), and restrSiteUtils (v1.2.9). HIV-1 LTR-containing fastq sequences were identified and filtered by allowing up to a maximum of five mismatches with the reference NL4-3 LTR sequence and if the LTR sequence had no match with any region of the human genome (GRCh37/hg19). Integration sites were determined from the sequence junction of the LTR and human genome sequences. All genomic sites in each dataset that hosted two or more sites (i.e. identical sites) were collapsed into one unique site for our analysis. Sites located in various common genomic features and non-B DNA motifs were quantified and heatmaps were generated using our in-house python program BLISIP Heatmap (BLISIPHA v1.0). Sites that could not be unambiguously mapped to a single region in the genome were excluded from the study. All non-B DNA motifs were defined according to previously established criteria^88^. Matched random control datasets were generated *in silico* as previously described^36^. Unique HIV LTRs were identified with BLISIP, aligned with MUSCLE^89^ and gap-stripped with trimAl^90^. All columns with gaps in more than 40% of the population were gap-stripped. Unique LTR sequence logos were generated using WebLogo^52^.

### Statistical Analyses

All statistical tests were performed as described in figure legends using GraphPad Prism 6.

### Data and software availability

The sequences reported in this paper will be deposited in the National Center for Biotechnology Information Sequence Read Archive (NCBI SRA).

## Author Contributions

M.-A.L. and S.D.B conceived the project and wrote the paper. T.M.R. performed the experiments. H.P.K. and M.D.C. performed the deep sequencing. H.O.A., T.M.R., M.-A.L. and S.D.B. performed the data analysis. All authors edited the manuscript.

## SUPPLEMENTARY INFORMATION

**Figure S1.**
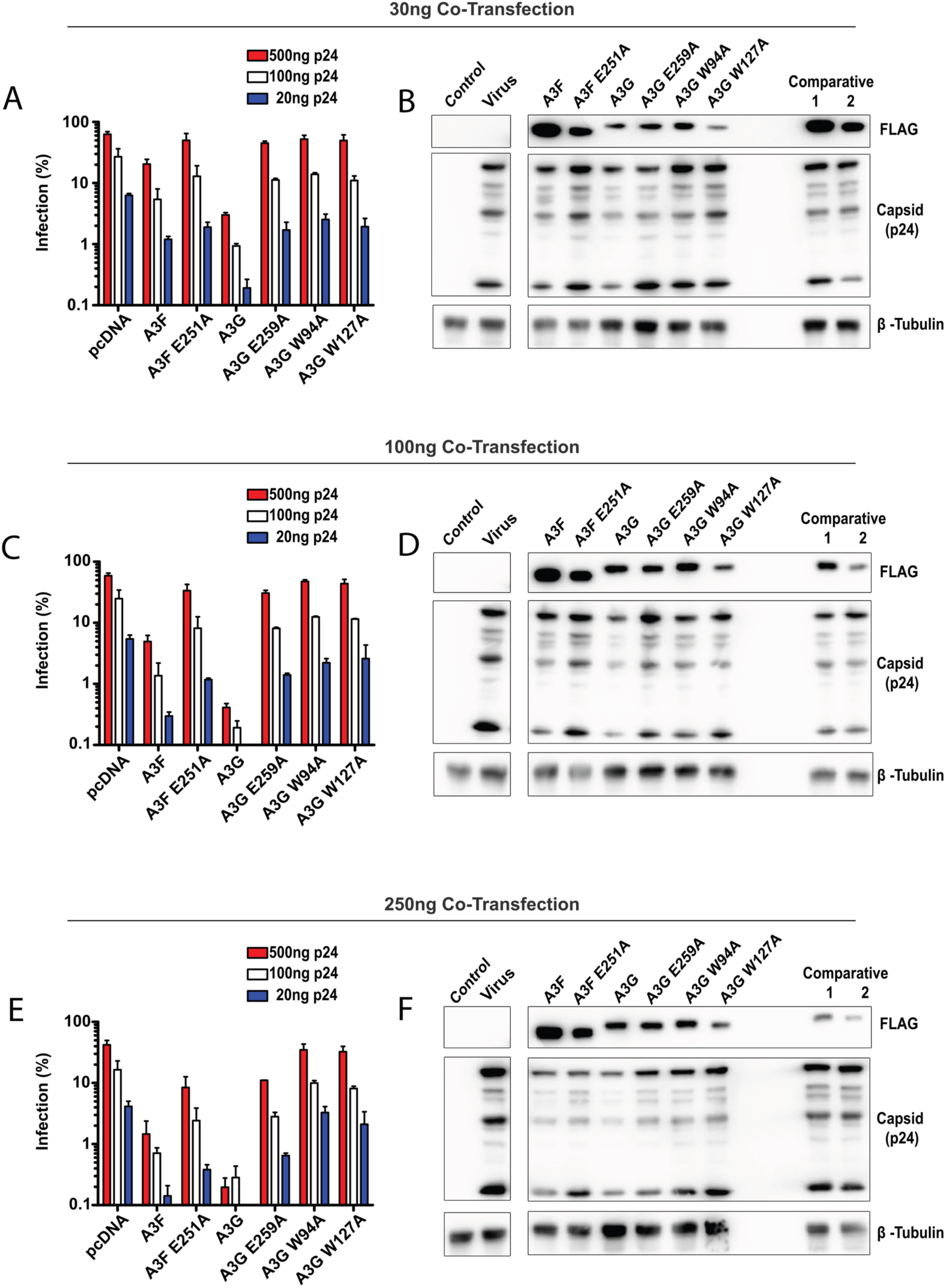
Effect of A3 proteins levels in virus producer cells on HIV-1 infection. VSV-G pseudotyped NL4-3-ΔEnv/ΔVif/eGFP was produced by co-transfection in 293T cells along with 30ng (**A** and **B**), 100ng (C and D) or 250ng (**E** and **F**) of A3 expression plasmids. Gel exposure controls, Comparative 1 and 2, represent co-transfection with 100ng of A3 W94A and A3 W127A plasmids, respectively. Control lane represents transfection with pcDNA3 only; virus lane is the co-transfection of pseudotyped virus along with the indicated amount (30ng, 100ng or 250ng) of pcDNA3 plasmid. CEM-SS cells were infected with pre-determined levels of capsid (p24) protein (20 ng, 100 ng or 500 ng) as measured by ELISA. The percentage of infected cells was determined by flow cytometry. Productively infected cells express the eGFP reporter gene encoded in the viral genome (**A, C** and **E**). Production of intracellular viral proteins was analyzed by Western blotting of transfected virus producer cells using anti-FLAG, anti-p24CA or anti-β tubulin antibodies (**B, D** and **F**).

**Figure S2.**
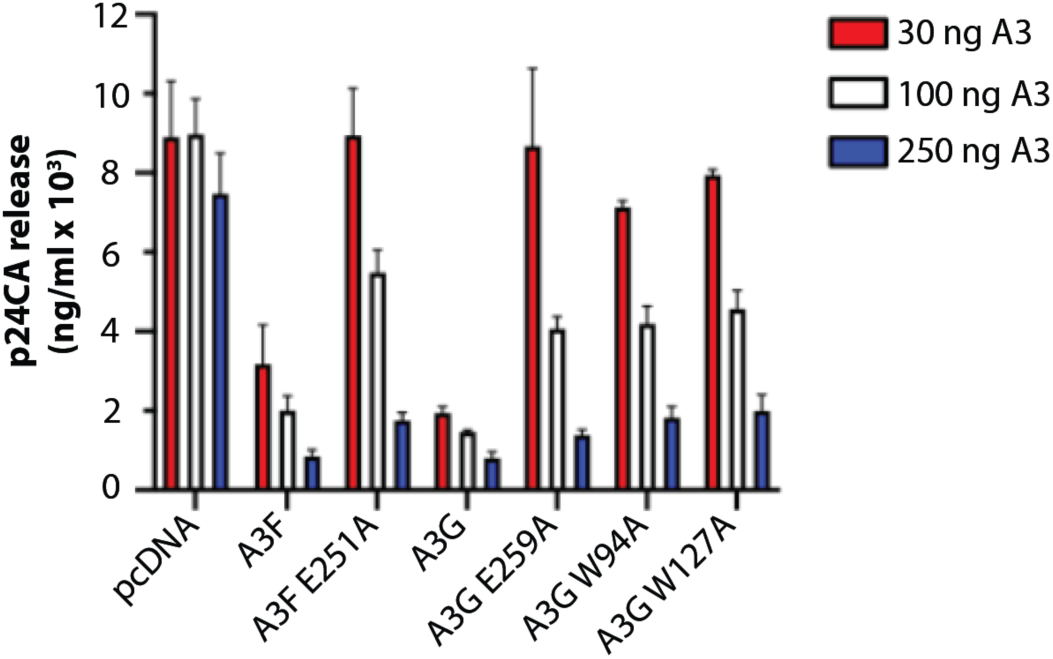
Measurements of HIV-1 particle release. Particle release was measured by p24CA sandwich ELISA for all transfection conditions (i.e., 30ng, 100ng or 250ng of co-transfected A3 or pcDNA3 plasmids).

**Figure S3.**
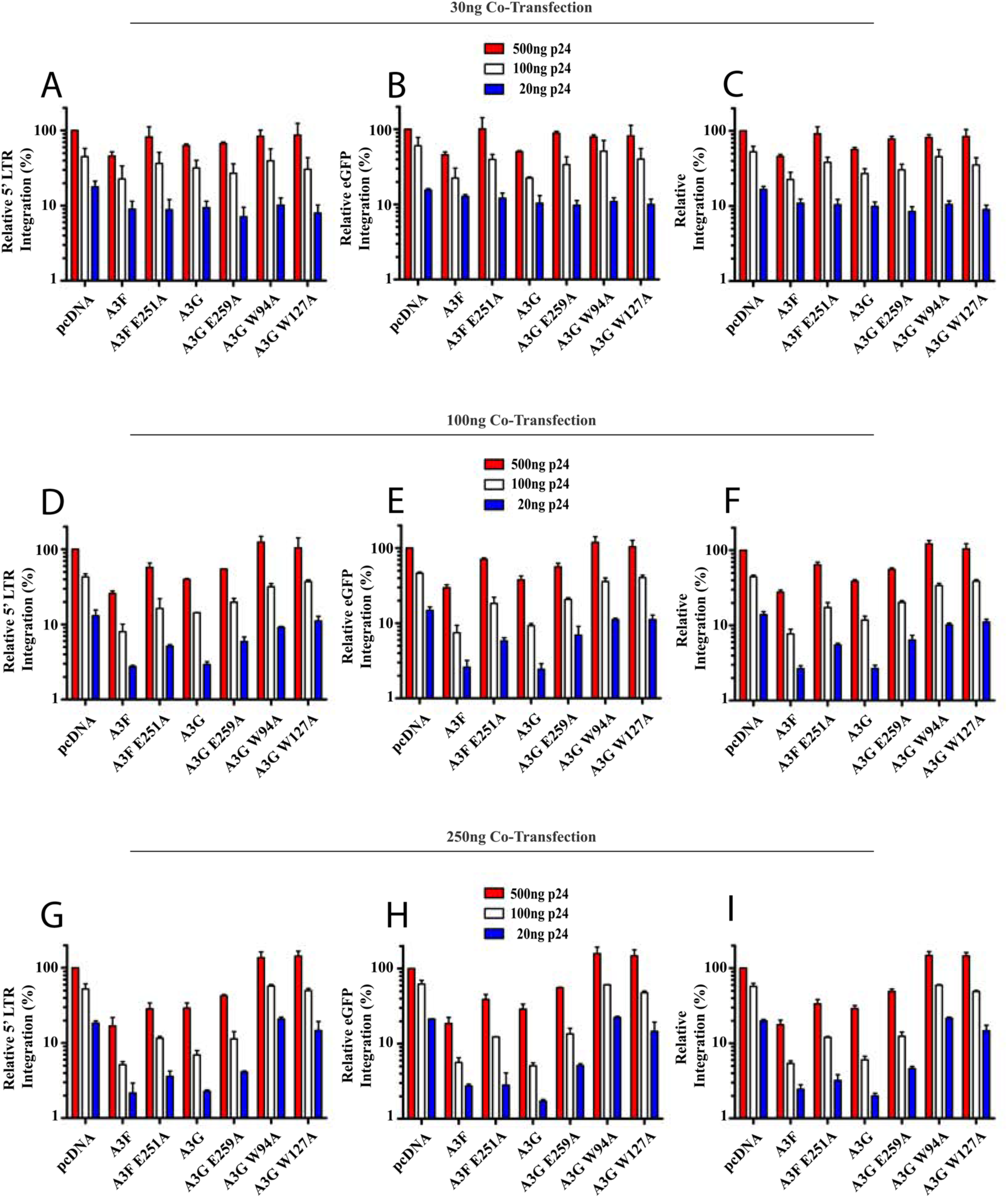
Effect of A3 protein levels in viral producer cells on HIV-1 integration. The level of integrated provirus in infected CEM-SS cells (as described in Figure S1) was determined by Alu-based ddPCR targeting either the 5’ LTR (**A, D** and **G**) or the *eGFP* reporter gene (**B, C** and **H**). Levels of proviral targets were normalized to a control gene (b-actin) by qPCR. Similar results were obtained for either integration analysis methods, with the average quantification of the two methods presented in panels **C, F** and **I**. Each panel is representative of the integration levels from cells transfected with 30 ng (**A-C**), 100 ng (**D-F**) and 250 ng (**G-I**) of A3 plasmid.

## SUPPLEMENTARY TABLE LEGENDS

**Table S1.** Integration site distribution in common genomic features from infected cells expressing various APOBEC3 constructs.

**Table S2.** Integration site heatmaps with fold changes in integration site abundance in and near common genomic DNA features from infected cells expressing various APOBEC3 constructs.

**Table S3.** Integration site distribution from HIV-1-infected CEM-SS cells expressing increasing concentrations of APOBEC3 or APOBEC3 mutant proteins.

**Table S4.** Integration site distribution in and near non-B DNA features from infected cells expressing various APOBEC3 constructs.

**Table S5.** Integration site heatmaps with fold changes in integration site abundance in and near non-B DNA features from infected cells expressing various APOBEC3 constructs.

**Table S6.** Effect of APOBEC3 expression on the number of integration hotspots.

**Table S7.** Integration site distribution in and near common genomic DNA and non-B DNA features of A3F-mutated LTRs.

**Table S8.** Integration site distribution in and near common genomic DNA and non-B DNA features of A3G-mutated LTRs.

